# Muscles of Facial Expression in Extinct Species of the Genus Homo

**DOI:** 10.1101/072884

**Authors:** Arturo Tozzi

## Abstract

We display a detailed description of mimetic muscles in extinct human species, framed in comparative and phylogenetic contexts. Using known facial landmarks, we assessed the arrangement of muscles of facial expression in *Homo sapiens, neanderthalensis, erectus, heidelbergensis* and *ergaster*. In modern humans, several perioral muscles are proportionally smaller in size (levator labii superioris, zygomaticus minor, zygomaticus major and triangularis) and/or located more medially (levator labii superioris, zygomaticus minor and quadratus labii inferioris) than in other human species. As mimetic musculature is examined in the most ancient specimens up to the most recent, there is a general trend towards an increase in size of corrugator supercillii and triangularis. *Homo ergaster’s* mimetic musculature closely resembles modern Homo, both in size and in location; furthermore, *Homo erectus* and *Homo neanderthalensis* share many muscular features. The extinct human species had an elaborate and highly graded facial communication system, but it remained qualitatively different from that reported in modern Homo. Compared with other human species, *Homo sapiens* clearly exhibits a lower degree of facial expression, possibly correlated with more sophisticated social behaviours and with enhanced speech capabilities. The presence of anatomical variation among species of the genus Homo raises important questions about the possible taxonomic value of mimetic muscles.

## INTRODUCTION

Primate muscles of facial expression (mimetic muscles) are unique in that they function either to open and close the apertures of the face or to tug the skin into intricate movements (Goodmurphy and Ovalle, 1999). The mimetic musculature is a visible signal of others’ social intentions and is used in transmitting close-proximity social information such as emotional states, individual recognition, mate, infant/caregiver interaction, promotion of social acceptance, moderation of the effects of social negative actions, territorial intentions and conflict of interests with strangers or competitors (Preuschoft, 2000; Burrows et al., 2006). The importance of the face as a critical variable in social intelligence is directly related to positive fitness consequences (Schmidt and Cohn, 2001).

Physical anthropologists have generally avoided the study of human facial expressions and nonverbal communication, leaving the interpretation largely to psychology and to other branches of anthropology (LaBarre, 1947; Birdwhistell, 1970). Although a number of studies have described facial muscular displays in Primate species (Pellatt, 1979; Swindler and Wood, 1982; Gibbs et al., 2002), results are not typically framed in an evolutionary perspective and remain focused on essentially nonadaptive questions. Furthermore, scarce data are available in ancient species of the genus Homo.

In our study, we compared the mimetic muscles of several human species, to provide some further phylogenetic perspective on the evolution of facial expression and its role in human social intelligence.

Using known facial landmarks, we assessed the arrangement of muscles of facial expression in modern *Homo sapiens* and in ancient human species (*Homo neanderthalensis*, *Homo erectus*, *Homo heidelbergensis* and *Homo ergaster*).

## MATERIALS AND METHODS

We evaluated the following cranial and/or mandibular specimens: modern African, Asian and European *Homo sapiens*; *Homo neanderthalensis* (La Chapelle-aux-Saints, Gibraltar 1, Amud 1, Kebara mandible, Krapina J mandible, Saccopastore 1, Shanidar 1);*Homo erectus* (skull of “Sinanthropus” from Zhoukoudian, new reconstruction after Tattersall and Sawyer 1996, Sangiran 17, Sangiran 1 mandible, PA 102 mandible, EV 9001); *Homo heidelbergensis* (Broken Hill 1, Petralona, Mauer mandible, AT 700, Arago 21), *Homo ergaster* (KNM-ER 3733, KNM-ER 992 mandible, KNM-ER 730 mandible, KNM-WT 15000).

Each sample was carefully assessed based on museum quality models made from the original fossils, on the latest literature and on life-size photographs, or some combination thereof.

The attachments of mimetic muscles were evaluated relative to known bony landmarks: the Frankfurt Horizontal, nasion, infra-orbital foramen, zygomaticomaxillary suture, zygomaticotemporal suture, maxillary incisive fossa, mandibular incisive fossa and mental foramen.

Following each measurement, we transferred the configuration of mimetic muscles to the image of a computer-manipulated skull of a model, thus evaluating the surface of muscular attachment. The skulls of different species were drawn so that their total vertical lengths were the same, allowing us to adjust in the same scale the measurements and the relationship distances.

Statistical analysis included Student’s t test and Fischer’s exact test. Significance was accepted at P<0.05. Results were expressed as mean±standard deviation.

### Muscles of facial expression in homo sapiens

We assessed in *Homo sapiens* the bony origin of the following muscles of facial expression (Stranding, 2004):

The corrugator supercilii muscle is attached to the inner extremity of the superciliary ridge. From the nasion, the mean point of origin is 4 mm lateral and 6 mm superior (Benedetto and Lahti, 2005).

The levator labii superioris muscle (quadratus labii superioris, infraorbital head) is attached to the inferior orbital margin immediately above the intraorbital foramen.

The zygomaticus minor muscle (quadratus labii superioris, zygomatic head) is attached to the malar surface of the zygomatic bone immediately behind the zygomaticomaxillary suture (Ferreira et al., 1997).

The zygomaticus major muscle is attached to the zygomatic bone, in front of the zygomaticotemporal suture (Spiegel and DeRosa, 2005). The upper extent of the lateral border of the zygomaticus major muscle is defined in relation to an oblique line extending from the mental protuberance to the notch at the junction of the frontal and temporal processes of the zygomatic bone. The lateral border of the zygomaticus major muscle is observed 4.4±2.2 mm lateral and parallel to this line (Mowlavi and Wilhelmi, 2004).

The caninus muscle (levator anguli oris) is attached to the canine fossa, below the orbital foramen.

The nasalis muscle, alar part (compressor naris, alar part) is attached to the incisive fossa and the incisive eminence of the central incisors.

The mentalis muscle (levator menti) is attached to the incisive eminence of the medial lower incisors.

The quadratus labii inferioris muscle (depressor labii inferioris) is attached to the external oblique line of the mandible, between the symphysis and the mental foramen.

The triangularis muscle (depressor anguli oris) is attached to the medial third of the external oblique line of the mandible.

Due to the lack of well-defined bony markings, the following muscles of facial expression were not evaluated, neither in modern nor in extinct human species: orbicularis oculi pars orbitalis, Horner’s muscle, levator labii superioris alaeque nasi, depressor septi nasi, nasalis (traverse part), incisivi labii superioris, buccinator, incisivi labii inferioris and platysma.

## RESULTS

The bony attachments of mimetic muscles in five human species are shown in **Figures 1** and **2**. The corrugator supercilii muscle of *Homo ergaster* and the mandibular mimetic muscles of *Homo heidelbergensis* were not evaluated, because of the lack of appraisable samples.

**Figure 1.**
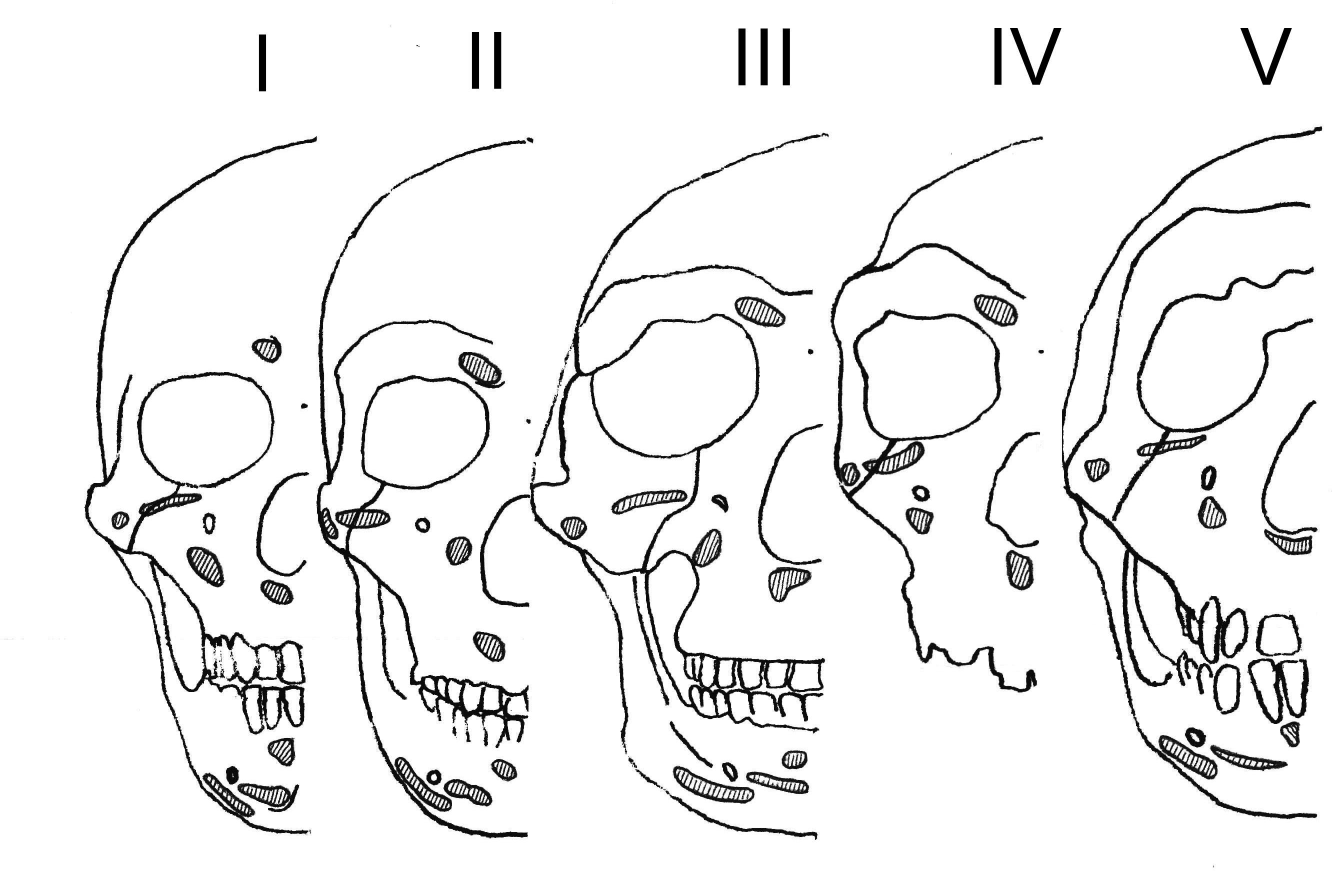
Schematical view illustrating the bony attachments of several mimetic muscles in five different human species. The skulls and the cranium are shown in frontal view. The skulls were drawn so that their total vertical lengths were the same. I:*Homo sapiens*; II:*Homo neanderthalensis*; III:*Homo erectus*; IV:*Homo heidelbergensis*; V:*Homo ergaster*.

**Figure 2.**
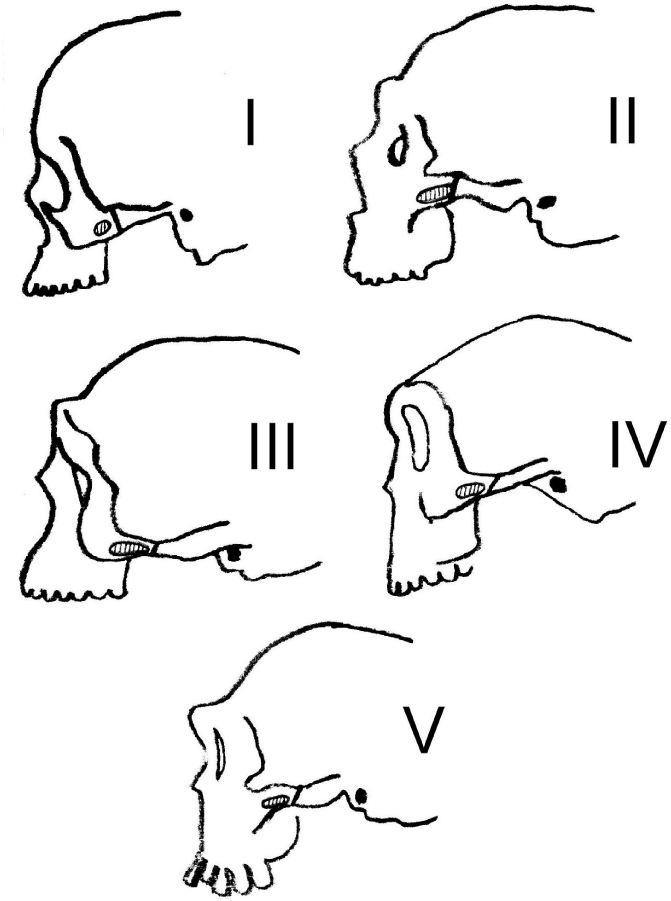
Schematical view illustrating the bony attachment of zygomaticus major muscle in five different human species. The crania are shown in lateral view. The crania were drawn so that their total vertical lengths were the same. I:*Homo sapiens*; II:*Homo neanderthalensis*; III:*Homo erectus*; IV:*Homo heidelbergensis*; V:*Homo ergaster*.

The mean surfaces of muscular attachments in different species of the genus Homo are shown in **Tables 1** and **2**. Four muscles (caninus, nasalis, mentalis and quadratus labii inferioris) did not show significant differences among human species and were not included in Tables.

**Table 1.**
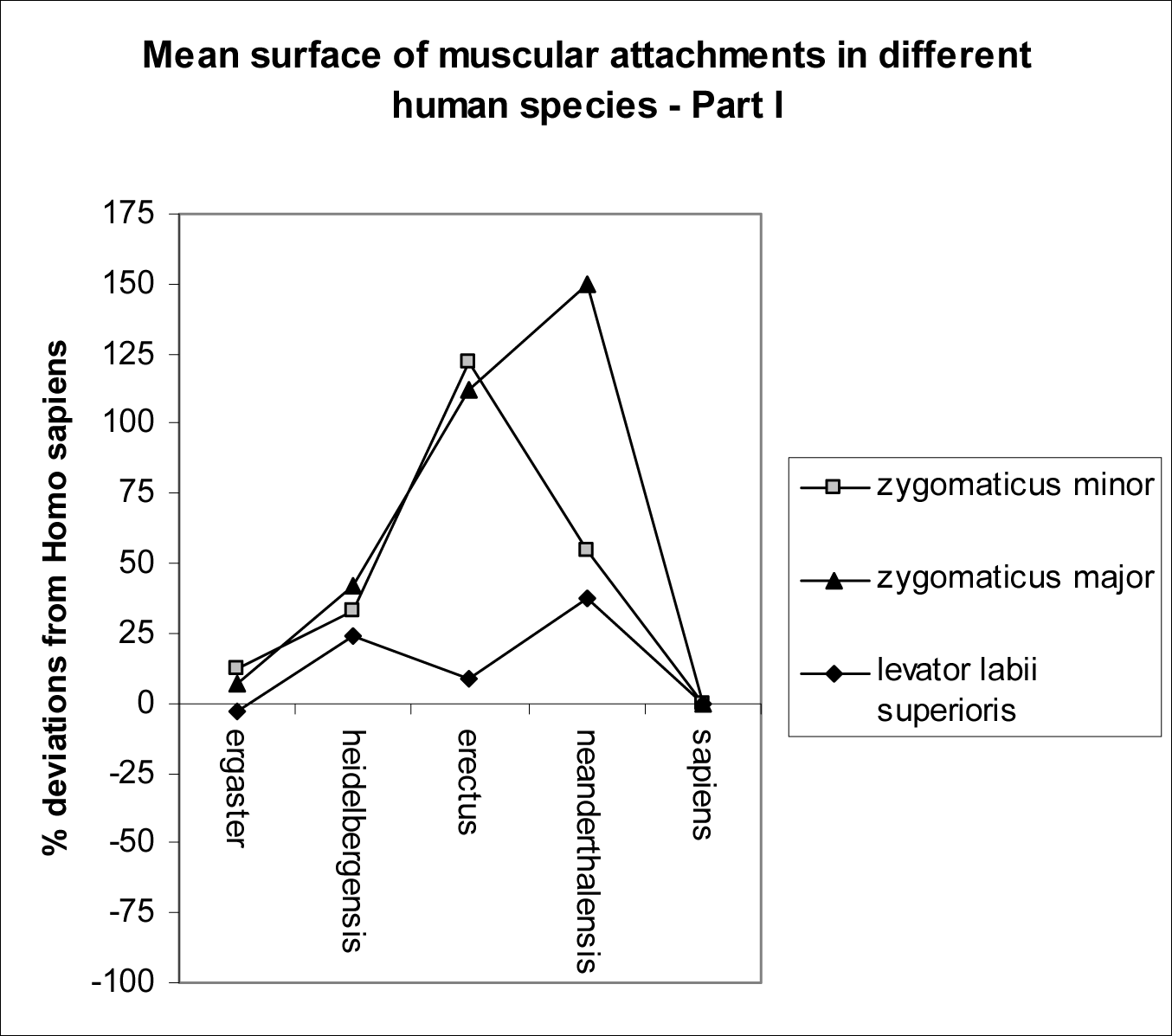
Mean surface of muscular attachments (zygomaticus major, zjgomaticus minor and levator labii superioris muscles) in different species of the genus Homo. Zero equals the *Homo sapiens* muscular surfaces; percent deviations from zero are the variation for extinct human species. All the values are statistically significant (p<0.001 or p<0.01, compared with *Homo sapiens*), apart from *Homo ergaster*’s values (p=N.S., compared with *Homo sapiens*).

**Table 2.**
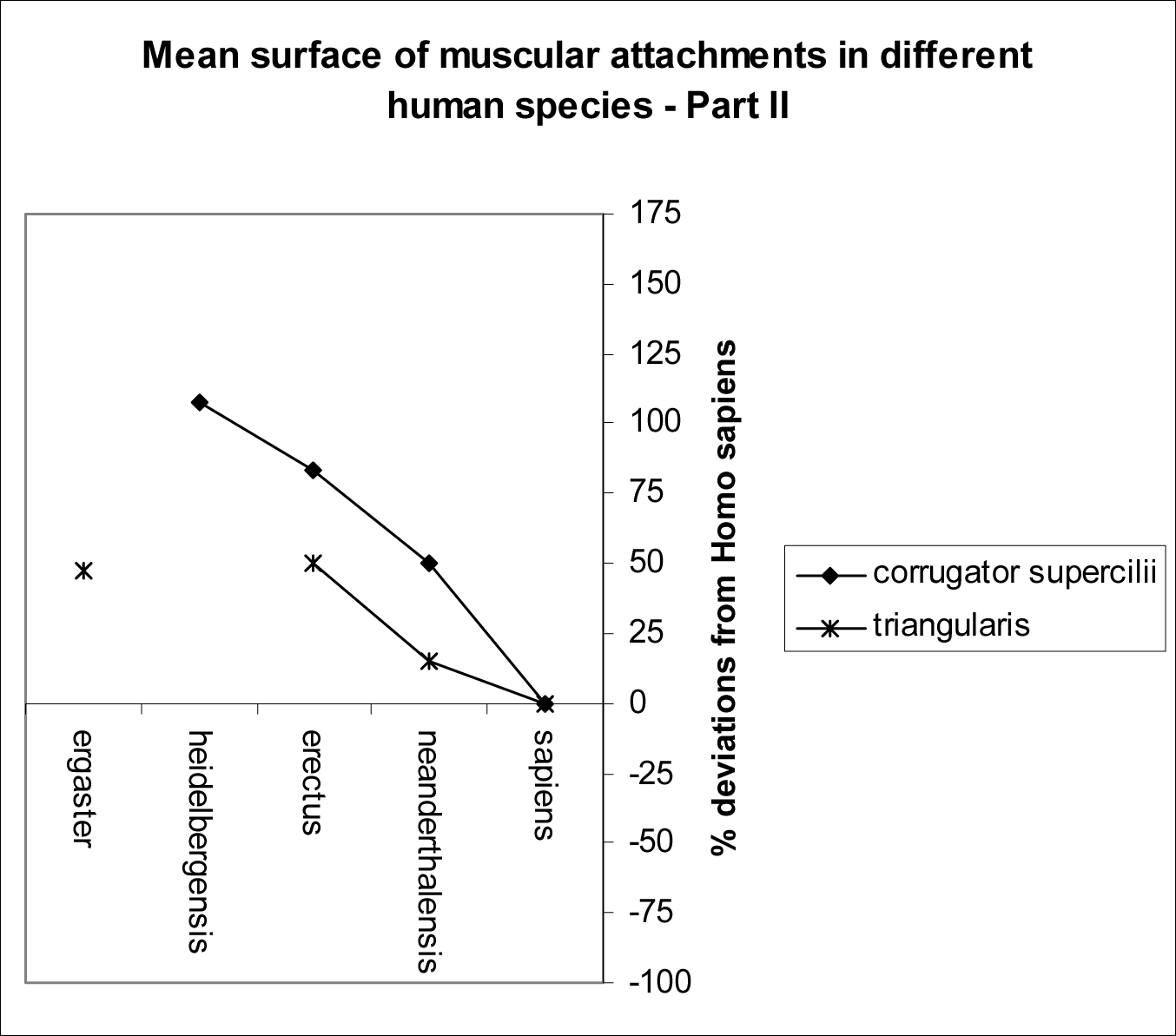
Mean surface of muscular attachments (corrugator supercilii and triangularis muscles) in different species of the genus Homo. Zero equals the *Homo sapiens* muscular surfaces; percent deviations from zero are the variation for extinct human species. All the values are statistically significant (p<0.001, compared with *Homo sapiens*).

Compared with extinct human species, *Homo sapiens* showed clear differences in mimetic muscles. In modern humans several perioral muscles were proportionally smaller in size (levator labii superioris, zygomaticus major, zygomaticus minor and triangularis) and/or located more medially (levator labii superioris, zygomaticus minor and quadratus labii inferioris), so that the muscular density of the oral region was higher than in ancient human species.

Furthermore, the mean surface of the corrugator supercilii was proportionally smaller in *Homo sapiens* than in other species of the genus Homo.

Compared with other human species, *Homo neanderthalensis* and *Homo erectus* shared the thickest and less crowded perioral musculature.

The mimetic muscules of *Homo ergaster* closely resembled modern human musculature, both in size and location.

## DISCUSSION

The current state of research in facial expression, combined with the topical interest in social intelligence as a driving force in human evolution, calls for the emergence of the study of mimetic muscles in physical anthropology (Schmidt and Cohn, 2001).

Although *Homo sapiens* has an elaborate repertoire of facial signals, there is very little research about the musculature underlying these movements, especially when compared with ancient human species. Here we present a detailed description of the mimetic muscles in species of the genus Homo, framed in comparative and phylogenetic contexts. By using this adaptive phenotype, it becomes possible to investigate facial expression in ancient humans from an evolutionary perspective (Fridlund, 1994).

The general belief is that the complexity of mimetic muscles increases from the most primitive Primates to the *Hominidae*, with the highest level of complexity being found in *Homo sapiens* (Gregory, 1929; Schultz, 1969). As species get more closely related to *Homo sapiens* and social networks become more intricate, it is held that their communicative facial repertoire and underlying facial musculature become more elaborate (Huber, 1930a, b; Preuschoft, 2000; Stranding, 2004). However, the validity of this hierarchical ascending phylogenetic model has been recently called into question. It has been found greater complexity in the facial muscles of *Otolemur* than previously reported (Burrows and Smith, 2003). Furthermore, it has been suggested that there is no foundation for claiming greater complexity in Homo facial expression musculature, compared with *Pan troglodytes* (Burrows et al., 2006). The present study showed that the mimetic muscles of extinct and modern human species are very similar; however, some important differences were found.

Compared with our ancient predecessors, mimetic muscles of modern Homo are thinner and more crowded. Many of the facial differences among human species concentrate on movements of the upper lip and midface: several muscles involved in disdain, sadness, smile and aggressive passions (corrugator supercilii, levator labii superioris, zygomaticus minor, zygomaticus major, triangularis) are less developed in *Homo sapiens*.

Zygomaticus major and zygomaticus minor muscles were strongly developed in *Homo erectus* and *Homo neanderthalensis*: the two muscles have high proportions of fast-twitch fibers relative to other muscles (Stal, 1994), indicating a possible specialization in these human species for fast facial movements.

Thus, the ancient species of the genus Homo had an elaborate and highly graded facial communication system, but it remained quantitatively and qualitatively different from that reported in *Homo sapiens*.

Human language is unique, and determining the time of origin would be vital in recognizing a key development in human history (Stringer and Andrews, 2005).

Among modern humans, social interaction almost invariably involves speech, with close relationships between facial expression and language. Facial expression is coordinated with speech at several levels: the use of mimetic muscles to articulate speech sounds (Massaro, 1998), the contribution of facial movements to the syntactic structure (Bavelas and Chovil, 1997) and the conversational signals that apply to the overall meaning of speech (Ekman, 1979).

Compared with *Homo neanderthalensis* and other extinct human species, *Homo sapiens* clearly exhibits a lower degree of facial expression, possibly correlated with enhanced speech capabilities and with more sophisticated social behaviours.

The link between decreased amount of facial expression and improved use of language could be the key to understand one of the *Homo sapiens*’ most sophisticated social abilities: the facial behaviours associated with deceit. Indeed, *Homo sapiens* is able to an effective suppression of facial expression in contexts where food resource, personal reputation or other positive outcome is at a stake (Ekman et al., 1997; Mitchell, 1999).

The presence of anatomical variation among human species raises important questions about the potential taxonomic value of mimetic muscles. Some Authors proposed that facial expression and correlated muscles of Primate taxa might be used in phylogenetic analyses (Gibbs et al., 2002; Burrows et al., 2006).

As mimetic musculature is examined in the most ancient specimens up to the most recent, there is a general trend towards an increase in size of corrugator supercillii and triangularis muscles.*Homo ergaster*’s mimetic muscles seem to have a closer relationship with *Homo sapiens*, while *Homo erectus* and *Homo neanderthalensis* share many muscular features. Further analyses are needed to elucidate the taxonomic value of our observations.

In conclusion, the findings from the present study provide some insight into the behavioural aspects of facial expression in extinct human species and their relationships with modern *Homo sapiens*.

## BIBLIOGRAPHY

1) Bavelas, J.B., Chovil, N., 1997. Faces in dialogue. In: Russell, J.A., Fernandez-Dols, J.M. (Eds.), The psychology of facial expression. Cambridge University Press, New York, pp. 334–346.

2) Benedetto, A.V., Lahti, J.G., 2005. Measurement of the anatomic position of the corrugator supercilii. Dermatol. Surg. 8 Pt 1, 923–927.

3) Birdwhistell, R.L., 1970. Kinesics and context. University of Pennsylvania Press, Philadelphia.

4) Burrows, A.M., Smith, T.D., 2003. Muscles of facial expression in Otolemur, with a comparison to Lemuroidea. Anat. Rec. 274A, 827–836.

5) Burrows, A.M., Waller, B.M., Parr, L.A., Bonar, C.J., 2006. Muscles of facial expression in the chimpanzee (Pan troglodytes): descriptive, comparative and phylogenetic contexts. J. Anat. 208, 153–167.

6) Ekman, P., 1979. About brows: emotional and conversational signals. In: von Cranach, M., Foppa, K., Lepenies, W., Ploog, D. (Eds.), Human ethology: claims and limits of a new discipline. Cambridge University Press, New York, pp. 169–222.

7) Ekman, P., Friesen, W.V., O’Sullivan, M., 1997. Smiles when lying. In: Ekman, P., Rosenberg, E. (Eds.), What the face reveals. Oxford University Press, New York, pp. 201–216.

8) Ferreira, L.M., Minami, E., Pereira, M.D., Chohfi, L.M., Andrews, J. de M., 1997. [Anatomical study of the levator labii superioris muscle] [Article in Portuguese] Rev. Assoc. Med. Bras. 43, 185–188.

9) Fridlund, A., 1994. Human facial expression: an evolutionary view. Academic Press, New York.

10) Gibbs, S., Collard, M., Wood, B., 2002. Soft-tissue anatomy of the extant hominoids: a review and phylogenetic analysis. J. Anat. 200, 3–49.

11) Goodmurphy, C.W., Ovalle, W.K., 1999. Morphological study of two human facial muscles: orbicularis oculi and corrugator supercilii. Clin. Anat. 12, 1–11.

12) Gregory, W.K., 1929. Our Face from Fish to Man. G.P. Putnam’s Sons, New York.

13) Huber, E., 1930a. Evolution of facial musculature and cutaneous field of trigeminus. Part I. Quar.t Rev. Biol. 5, 133–188.

14) Huber, E., 1930b. Evolution of facial musculature and cutaneous field of trigeminus. Part II. Quar.t Rev. Biol. 5, 389–437.

15) LaBarre, W., 1947. The cultural basis of emotions and gestures. J. Pers. 16, 49–68.

16) Massaro, D.W., 1998. Perceiving talking faces. MIT Press, Cambridge.

17) Mitchell, R.W., 1999. Deception and concealment as strategic script violation in great apes and humans. In: Parker, S.T., Mitchell, R.W., Miles, H.L. (Eds.), The mentalities of gorillas and orangutans. Cambridge University Press, New York, pp. 295–315.

18) Mowlavi, A., Wilhelmi, B.J., 2004. The extended SMAS facelift: identifying the lateral zygomaticus major muscle border using bony anatomic landmarks. Ann. Plast. Surg. 52, 353–357.

19) Pellatt, A., 1979. The facial muscles of three African primates contrasted with those of Papio ursinus. S. Afr. J. Sci. 75, 436–440.

20) Preuschoft, S., 2000. Primate faces and facial expressions. Soc. Res. 67, 245–271.

21) Schmidt, K.L., Cohn, J.F., 2001. Human Facial Expressions as Adaptations: Evolutionary Questions in Facial Expression Research. Yearb. Phys. Anthropol. 44, 3–24.

22) Schultz, A.H., 1969. The Life of Primates. Universe Books, New York.

23) Spiegel, J.H., DeRosa, J., 2005. The anatomical relationship between the orbicularis oculi muscle and the levator labii superioris and zygomaticus muscle complexes. J. Plast. Reconstr. Surg. 116, 1937–1942.

24) Stal, P., 1994. Characterization of human oro-facial and masticatory muscles with respect to fibre types, myosins and capillaries. Morphological, enzyme-histochemical, immuno-histochemical and biochemical investigations. Swed. Dent. J. Suppl. 98, 1–55.

25) Stranding, S., 2004. Gray’s Anatomy, 39th edn. Churchill Livingstone, London.

26) Stringer, C., Andrews, P., 2005. The complete world of Human evolution. Thames & Hudson, London.

27) Swindler, D.R., Wood, C.D., 1982. An Atlas of Primate Gross Anatomy. Malabar, F.L. Robert E. Krieger Publishing.

28) Tattersall, I.; Sawyer, G.J., 1996. The skull of “Sinanthropus” from Zhoukoudian, China: a new reconstruction. J. Hum. Evol., 31, 311–314.

